# Convergence, plasticity, and tissue residence of regulatory T cell response via TCR repertoire prism

**DOI:** 10.1101/2023.06.15.544726

**Authors:** T.O. Nakonechnaya, B. Moltedo, E.V. Putintseva, S. Leyn, D.A. Bolotin, O.V. Britanova, M. Shugay, D.M. Chudakov

**Author notes:** These authors contributed equally (T.O. Nakonechnaya, B. Moltedo, E.V. Putintseva).

## Abstract

Suppressive function of regulatory T cells (Treg) is dependent on signaling of their antigen receptors triggered by cognate self, dietary or microbial antigens in the form of peptide-MHC class II complexes. However, it remains largely unknown whether distinct or shared repertoires of Treg TCRs are mobilized in response to different challenges in the same tissue or the same challenge in different tissues. Here we used a fixed TCRβ chain FoxP3-GFP mouse model to analyze conventional (eCD4) and regulatory (eTreg) effector TCRα repertoires in response to six distinct antigenic challenges to the lung and skin. This model showed highly “digital” repertoire behavior, allowing for easy-to-track challenge-specific TCRα CDR3 clusters. For both studied subsets, we observed challenge-specific clonal expansion yielding homologous TCRα clusters within and across animals and exposure sites, which were also reflected in the draining lymph nodes but not systemically. Some clusters were shared across cancer challenges, suggesting response to common tumor-associated antigens. For most challenges, eCD4 and eTreg clonal response did not overlap, indicating the distinct origin of the two cell subsets. At the same time, we observed such overlap at the sites of certain tumor challenges. The overlaps included dominant responding TCRα motif and characteristic iNKT TCRα, suggesting the tumor-induced eCD4-eTreg plasticity. Additionally, our TCRα repertoire analysis demonstrated that distinct antigenic specificities are characteristic for eTreg cells residing in particular lymphatic tissues, regardless of the challenge, revealing the homing-specific, antigen-specific resident Treg populations. Altogether, our study highlights both challenge-specific and tissue-specific responses of Treg cells associated with distinct clonal expansions.

**Graphical abstract:** 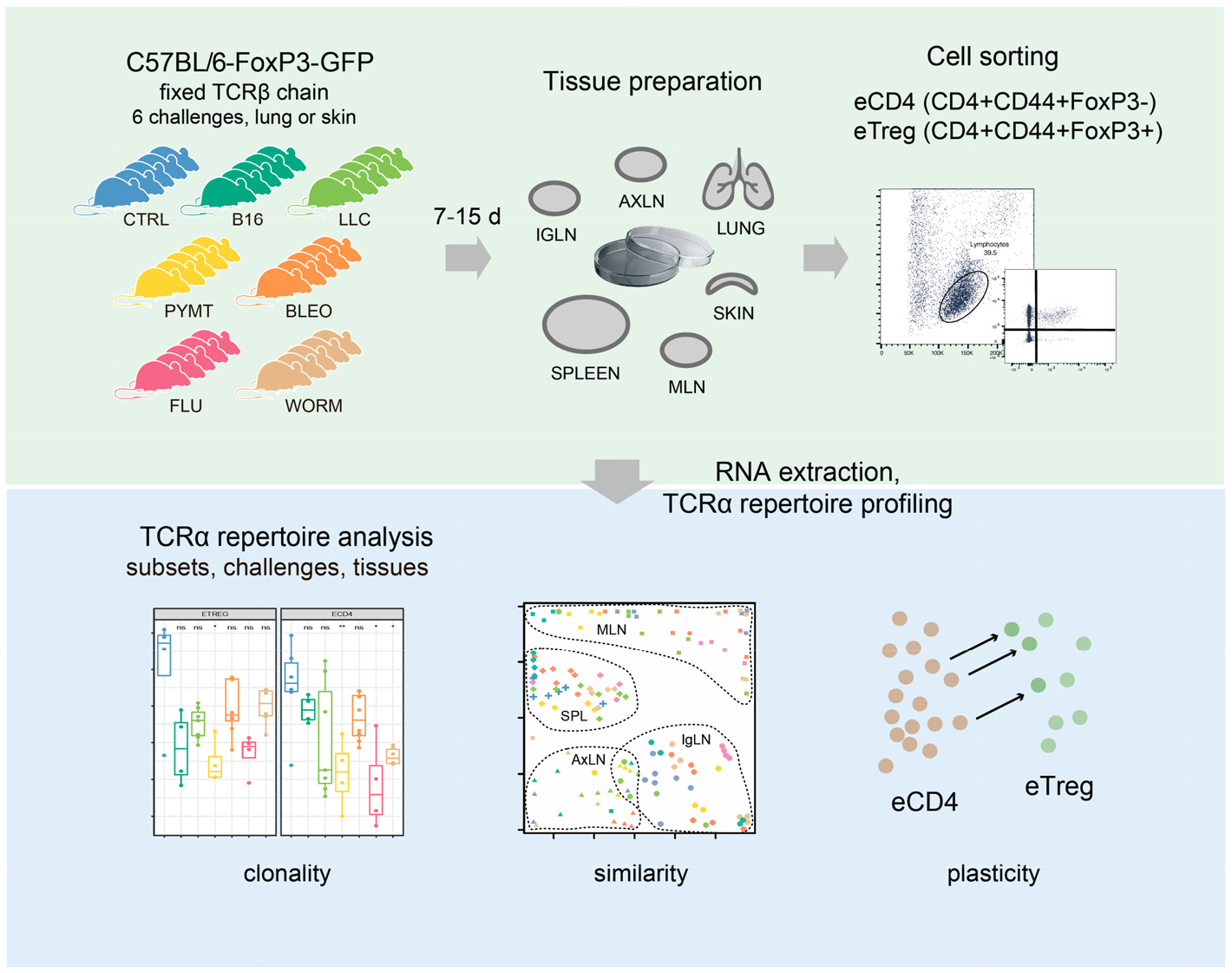

## Introduction

Regulatory T cells (Treg), which constitute 5-10% of peripheral CD4^+^ T cells are indispensable for the maintenance of immunological self-tolerance^1,2^ and regulation of ongoing immune responses to microbiotal^3-5^ and foreign^6-9^ antigens, contributing to tissue and metabolic homeostasis and regeneration^2^. Treg suppress immune responses through various mechanisms, but primarily do so via antigen-specific^10^ interaction with professional and non-professional antigen-presenting cells^11^. There is considerable data suggesting that Treg employ higher-affinity T cell receptor (TCRs) and CTLA4 to compete with conventional CD4^+^ T cells in an antigen-specific manner^12^.

Most Treg cells are produced by the thymus as a functionally distinct FoxP3^+^ T cell subset. Thymic Treg are generated based on positive selection against self-peptide/MHCII (pMHCII) complexes in the thymic medulla, and multiple other immune cell types may further contribute to this process, including medullary thymic epithelial cells (mTECs), resident and immigrating dendritic cells, macrophages, and B cells^2,13-15^.

Here we investigated the TCR landscape of conventional (eCD4) and regulatory (eTreg) effector CD4^+^ T cells in lungs, spleen, skin, and three lymph node locations after response to six distinct antigenic challenges to the lungs or skin (see **Graphical abstract**). We used FoxP3-GFP mice with a fixed TCRβ context^16^ and analyzed the TCRα repertoire composition for eCD4 and eTreg cells.

The adaptive immune system in this mouse model relies entirely on TCRα diversity to generate antigen-specific responses. On one hand, the relatively limited TCRα repertoire^16^ results in highly convergent responses that are easily traceable across groups of mice facing the same challenge. The selection of the same or highly similar TCRα sequences against each antigen leads to the formation of clusters of homologous TCRα CDR3s within and between mice. On the other hand, this model allows for the tracking of all challenge-specific or tissue-specific features of the TCR repertoire using only one chain^16^. Overall, compared to conventional mice, this model provides a more powerful means of monitoring convergent TCR responses. At the same time, the model generally mimics the natural behavior of the full-fledged TCRαβ T cell repertoire.

Both for eCD4 and eTreg subpopulations, our analysis of the TCRα repertoire on the fixed TCRβ background revealed a highly focused response which was distinct for each of the antigenic challenges. The response was clonotypically shared between individual mice and between different tissues and formed clear convergent^17^ TCRα CDR3 sequence motifs responding to the challenges.

In conditions with such a highly convergent TCR response involving multiple independently-primed T cell clones with similar TCRα CDR3 sequences, one could expect a similar repertoire of clonotypes to dominate within the eCD4 and eTreg subpopulations, given the assumption that both subsets may start from the same pool of antigen-inexperienced naïve T cells. However, we demonstrate that the TCR landscape of eCD4 and eTreg responses was distinct, suggesting that these effector cell types originate from distinct T cell subsets.

The exceptions were the antigenic challenges with Lewis lung carcinoma (LLC) and MMTV-PyMT-derived tumor cells (PYMT), where we observed increased overlap between the eCD4 and eTreg repertoires at the site of challenge, but not systemically. A dominant eCD4 TCRα motif was also present in corresponding eTreg subsets in the LLC and PYMT challenge.

Similarly, the characteristic innate natural killer T cell (iNKT) TCRα variants (that were abundant within eCD4 subsets upon challenge with LLC, bleomycin, and the helminth *Nippostrongylus brasiliensis*) were represented in the corresponding eTreg subset in the LLC challenge experiments but not with the bleomycin and helminth antigens. We attribute these observations to the transient plasticity of effector CD4^+^ T cells in the context of an immunosuppressive tumor burden.

Furthermore, we reveal tissue-specific eTreg TCRα CDR3 motifs that were always present in specific tissue locations irrespectively of the applied antigenic challenge. This observation highlights the existence of the diverse homing-specific, antigen-specific resident Treg populations.

Altogether, our results show the highly antigen-specific and distinct eCD4 and eTreg responses to different challenges, supporting the initial thymic programming of Treg cells responding to newly incoming challenges, and suggest that the fixed TCRβ chain, FoxP3-GFP mouse offers a valuable “digital” model of TCR response that benefits from highly convergent, easy-to-track TCRα CDR3 motifs.

## Results

### TCRα repertoire sequencing

Six distinct challenges including influenza virus, *N. brasiliensis*, bleomycin-induced injury, LLC, PYMT, and B16 melanoma were applied to the lung and skin of *Foxp3*^*gfp*^ *Tcra*^*−/+*^ mice bearing the DO11.10 TCRβ transgene^18^ (3–7 mice per group, see **Supplementary Table 1** for details on each tissue and challenge experiment). After incubation, mice were sacrificed and tissue from the lungs, spleen, skin, and three types of lymph nodes (lung-draining mediastinal, MLN, axillary, AXLN, and intraglandular, IGLN) were isolated and digested to generate single-cell suspensions. eCD4 and eTreg CD4^+^ cells were sorted based on FoxP3-GFP signal. RNA-based unique molecular identifier (UMI)-labeled TCRα cDNA libraries were obtained using a previously-reported technique^19,20^ with minor modifications, and then analyzed using MIGEC^21^ and MiXCR^22^ software. See **Graphical abstract** for the experimental scheme. Altogether, we sequenced 524 TCRα cDNA libraries, yielding 13,605 ± 14,948 UMI-labeled TCRα cDNA molecules and 3,392 ± 3,076 TCRα CDR3 clonotypes per eTreg sample, and 37,412 ± 33,129 UMI-labeled TCRα cDNA molecules and 5,518 ± 4,601 TCRα CDR3 clonotypes per eCD4 sample (see **Supplementary Table 2** for details on each cloneset).

Lungs are exposed to a myriad of different insults during their lifetime and are constantly in need of a T cell response to resolve inflammatory insults. Furthermore, Treg with an effector phenotype accumulate in the inflamed lung^23^ but it remains poorly understood whether there is a common antigenic denominator driving TCR specificities in the tissue or whether there are selective TCR subsets expanded upon a particular inflammatory insult. Therefore, in the following analyses, we mainly focused on TCR repertoires obtained from T cells infiltrating the lung tissue, for which we have also obtained the largest collection of samples, with several more specific cross-tissue analyses showing the similarity of systemic T cell responses.

### Clonality of eCD4 and eTreg repertoires in the lung

First, we explored how clonal the eCD4 and eTreg response is in lungs following distinct antigenic challenges. TCRα repertoire diversity was assessed using several widely used metrics, including observed diversity (number of distinct clonotypes), normalized Shannon Wiener index (repertoire evenness and the extent of clonal expansion) and Chao 1 (estimated lower bound of total diversity based on relative representation of small clonotypes). For normalization, all of these metrics were obtained from datasets that had been down-sampled to 1,000 randomly-chosen, UMI-labeled TCRα cDNAs^19^. We observed a prominent clonal response to each of the challenges compared to the control group (**Fig. 1a–c**). The decrease in diversity metrics was comparable for the lung-infiltrating eCD4 and eTreg cells. This indicates that the amplitude and focused nature of the effector and regulatory T cell response in lungs is generally comparable.

**Figure 1.**
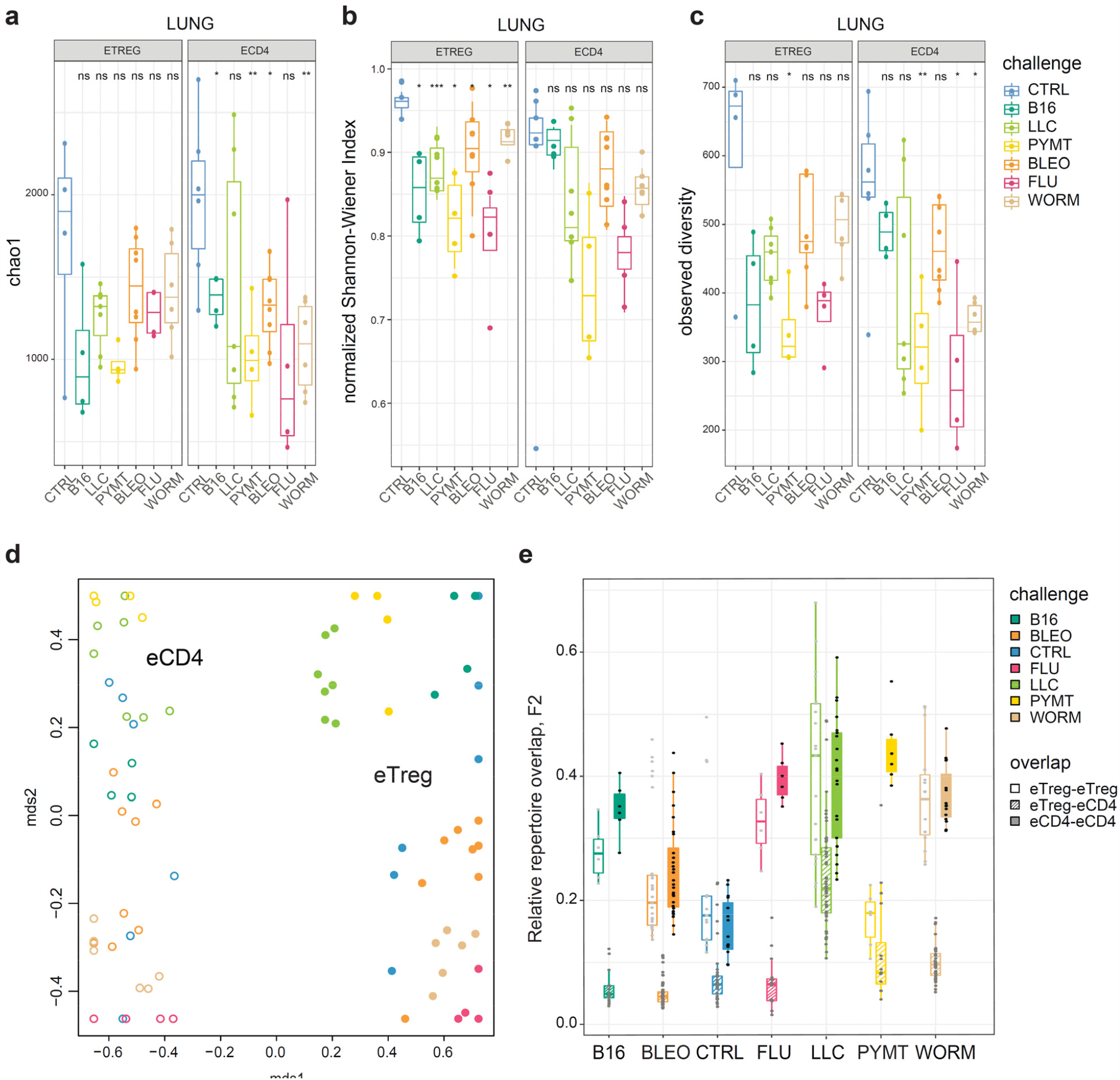
eTreg and eCD4 subsets. **a-c**. Clonality and diversity of lung eTreg and eCD4 subsets in response to distinct antigenic challenges. We calculated the (**a**) Chao1 estimator, (**b**) normalized Shannon-Wiener index, and (**c**) observed diversity for each TCRα repertoire obtained from each CD4^+^ T-cell subset from each animal (3 < n < 7) with different antigenic challenges. P-values shown as: ^*^p < 0.05, ^**^p < 0.01, ^***^p < 0.001, and ^****^p < 0.0001, based on parametric t-test for each group versus control. **d**. Relative overlap between amino acid-defined lung TCRα CDR3 repertoires, visualized as a VDJtools MDS plot. Euclidean distance between points reflects the distance between repertoires. Data were normalized to the top 1,000 most frequent clonotypes and weighted by clonotype frequency (F2 metric in VDJtools). Clonotypes were matched on the basis of identical TRAV gene segments and identical TCRα CDR3 sequences. The closer two circles are, the higher the overall frequency of shared clonotypes. Node colors correspond to the challenge antigen. Filled circles are lung eTreg, open circles are lung eCD4 cells. **e**. The same F2 metrics as in (**d**), shown separately for each challenge and for eTreg-eTreg, eTreg-eCD4, and eCD4-eCD4 repertoire overlap. For both (**d**) and (**e**), mice were analyzed in an all-versus-all fashion, irrespective of whether eTreg and eCD4 subsets were obtained from the same or distinct mice.

### Responding eCD4 and eTreg repertoires are distinct

Next, we analyzed all-versus-all pairwise overlap of the amino acid TCRα CDR3 repertoires of lung-infiltrating eTreg and eCD4 cells. We used F2 similarity metrics in the VDJtools software^24^, which employs a clonotype-wise sum of geometric mean frequencies that takes into account the relative size of shared clonotypes. As such, F2 metrics generally enable comparison of pairs of TCR repertoires with regard to the relative share occupied by common clonotypes. This analysis showed that the eTreg and eCD4 repertoires are highly distinct across distinct challenges (**Fig. 1d,e**).

### eTreg response is convergent and differs for distinct challenges

We next zoomed in on the eTreg repertoire in order to investigate how focused their antigen-specific response is in the lungs. Repertoire overlap analysis showed that eTreg repertoires were highly similar across mice for each of the challenges, and differed between challenges (**Fig. 2a, b**). Cluster sequence analysis further revealed common dominant TCRα CDR3 motifs in response to each of the challenges (**Fig. 2a, Supplementary Figs. 1,2, Supplementary Tables 4**,**5**). It should be noted that the eTreg repertoires in the three different cancer challenges (B16, LLC, and PYMT) clustered together and shared some CDR3 motifs, suggesting response to shared tumor-associated antigens. In LLC and PYMT, but not in other challenges, some of the characteristic CDR3 motifs were shared with the eCD4 subset.

**Figure 2.**
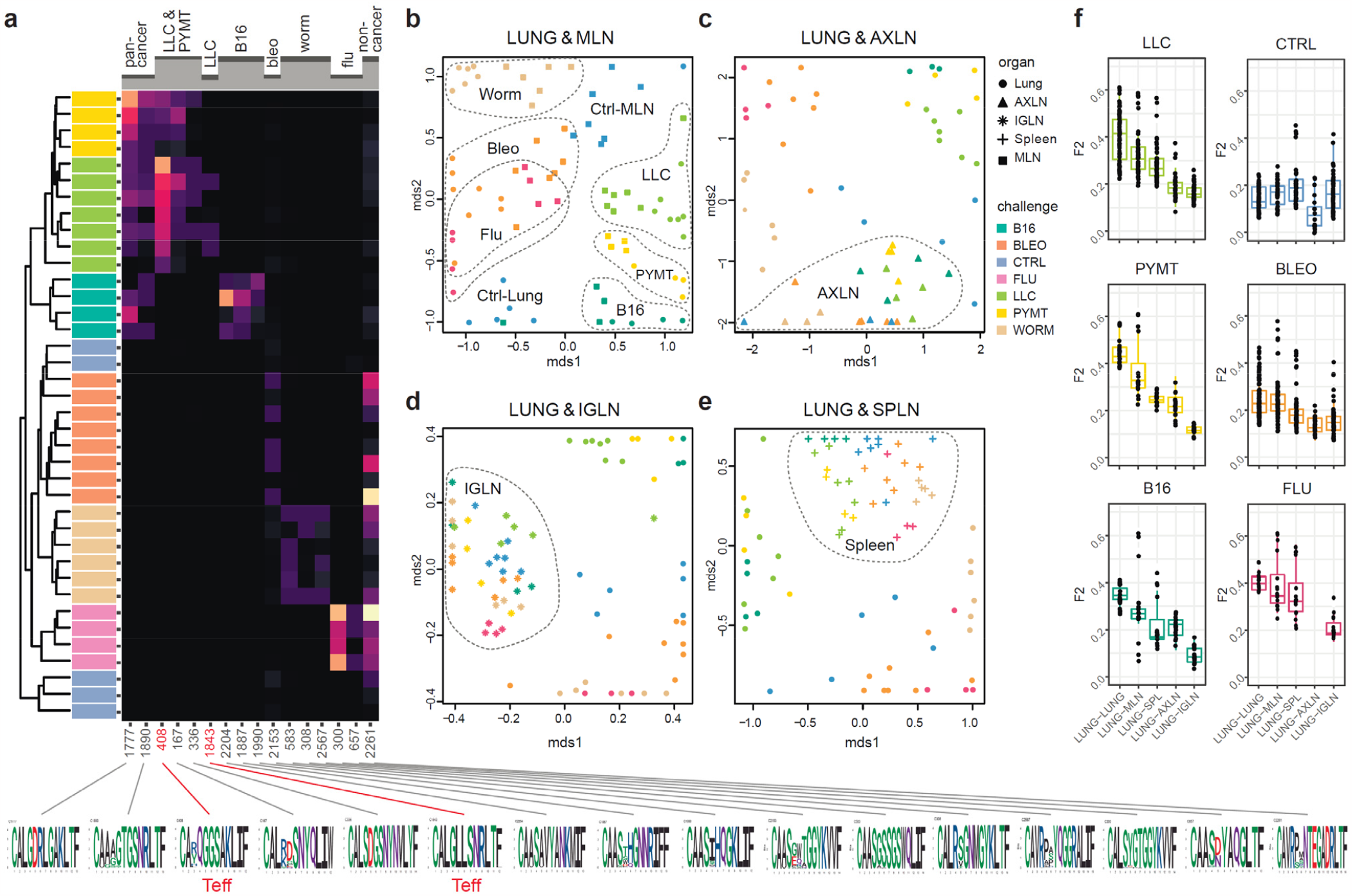
eTreg TCR repertoire in various tissues following antigen challenge in the lung. Relative overlap between amino acid-defined eTreg TCRα CDR3 repertoires visualized using VDJtools. Data were normalized to the top 1,000 most frequent clonotypes and weighted by clonotype frequency (F2 metric). Clonotypes were matched on the basis of identical TRAV gene segments and identical TCRα CDR3 sequences. (**a**) Dendrogram of Lung eTreg TCRα CDR3 repertoires, where branch length shows the distance between repertoires. Heat map at right shows the distribution of selected TCRα CDR3 clusters associated with specific challenges. Corresponding TCRα CDR3 logos are shown at bottom; clusters also observed in corresponding eCD4 samples are indicated in red. (**b**–**e**) MDS plots comparing repertoires between pairs of tissues, where the Euclidean distance between points reflects the distance between repertoires. Overlap is shown for (**b**) lung versus MLN, (**c**) lung versus AXLN, (**d**) lung versus IGLN, and (**e**) lung versus spleen. The closer two samples are, the higher the overall frequency of shared clonotypes. Node colors correspond to different challenges. Mice were analyzed in an all-versus-all fashion, irrespective of whether tissues were obtained from the same or distinct mice. Circles: lung. Squares: MLN. Triangles: AXLN. Snowflakes: IGLN. Crosses: spleen. (**f**) Graphs show eTreg F2 repertoire overlap between lung tissue from different animals, and between lungs and other tissues of all animals.

### eTreg repertoire upon lung challenge is reflected in the draining lymph node

It has previously been shown in a mouse model that the antigen-specific Treg response to influenza in the lungs is also reflected in the lung-draining mediastinal lymph node (MLN)^25^. Our data revealed this at the repertoire level, showing that the same TCR clonotypes and CDR3 motifs distinguish antigen-specific eTreg response to each of the different lung challenges in the lung and MLN (**Fig. 2b, Supplementary Figs. 1,2, Supplementary Tables 4**,**5**). It should be noted that each challenge produces its own specific response, wherein eTreg TCRα CDR3 repertoires from the lung and MLN are located side by side. At the same time, the lung and MLN eTreg repertoires of control mice obtained in the absence of any antigenic challenges were distinct, confirming that repertoire similarity is dictated by corresponding antigenic challenges (**Fig. 2b**). In contrast, the eTreg repertoires in distant AXLN and IGLN lymph nodes and the spleen were less similar to that observed in the lungs, and clustered separately (**Fig. 2c–f**). Altogether, these results demonstrate the selective tissue localization of the antigen-focused Treg response.

### Repertoire focus is the same for the lung and skin tumor localization

The eTreg TCR repertoire in lungs (upon lung tumor challenge) and in skin (upon corresponding tumor challenge in skin) was also highly similar (**Fig. 3**), demonstrating that the antigen-specific character of the Treg response dominates over the tissue location of the challenge. TCR repertoires from eCD4 T cells in the lungs and MLN in response to distinct lung challenges, as well as repertoires of skin eCD4 T cells in response to various skin challenges also clustered together (**Fig. 3**), with a magnitude of repertoire convergence that was at the same level as for eTreg cells (**Fig. 1e**).

**Figure 3.**
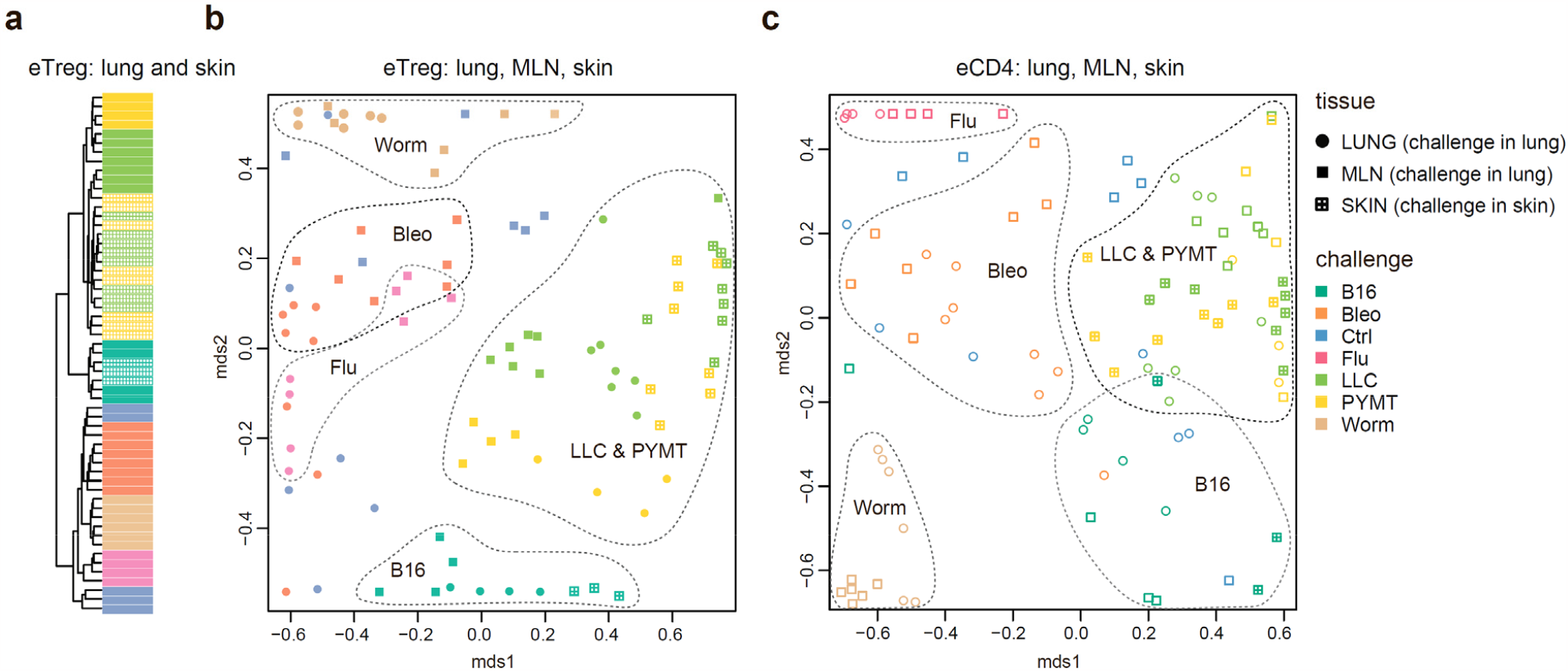
Convergence of eTreg and eCD4 TCR response in distinct tissues. (**a**,**b**) Relative overlap between amino acid-defined lung and MLN (upon lung challenge) and skin (upon skin challenge) eTreg TCRα CDR3 repertoires as visualized using VDJtools. Data were normalized to the top 1,000 most frequent clonotypes and weighted by clonotype frequency (F2 metric in VDJtools). Clonotypes were matched on the basis of identical TRAV gene segments and identical TCRα CDR3 sequences. (**a**) Dendrogram branch length shows the distance between repertoires. (**b**) Euclidean distance between points on the MDS plot reflects the distance between repertoires. The closer two samples are, the higher the overall frequency of shared clonotypes. Mice were analyzed in an all-versus-all fashion, irrespective of whether tissues were obtained from the same or distinct mice. Node colors correspond to different challenges. Circles: lung. Crossed squares: skin. (**c**) Relative overlap between amino acid-defined lung and MLN (upon lung challenge) and skin (upon skin challenge) eCD4 TCRα CDR3 repertoires.

### eCD4 to eTreg conversion is only observed in two cancer challenges

As can be seen in **Figure 1e**, TCRα CDR3 eTreg and eCD4 amino acid repertoires in the lungs were more similar in the context of the LLC tumor challenge compared to other challenges, and this could reflect induced plasticity of the eCD4 subset. In order to assess possible eCD4-to-eTreg clonal conversion due to natural helper T cell plasticity, we analyzed eTreg versus eCD4 repertoire overlap at the level of CDR3a nucleotide clonotypes for paired eCD4-eTreg samples obtained from the same mice. Estimated eCD4-to-eTreg conversion was strongest in skin and in lung following LLC and PYMT challenge, but was low or absent in the aftermath of other antigenic challenges, including B16 tumor (**Fig. 4a,c**). The effect was probably local and transient, and was not observed at systemic level in the spleen (**Fig. 4b,d**). CDR3a cluster #408 was the major contributor to the observed conversion (**Fig. 4e**).

**Figure 4.**
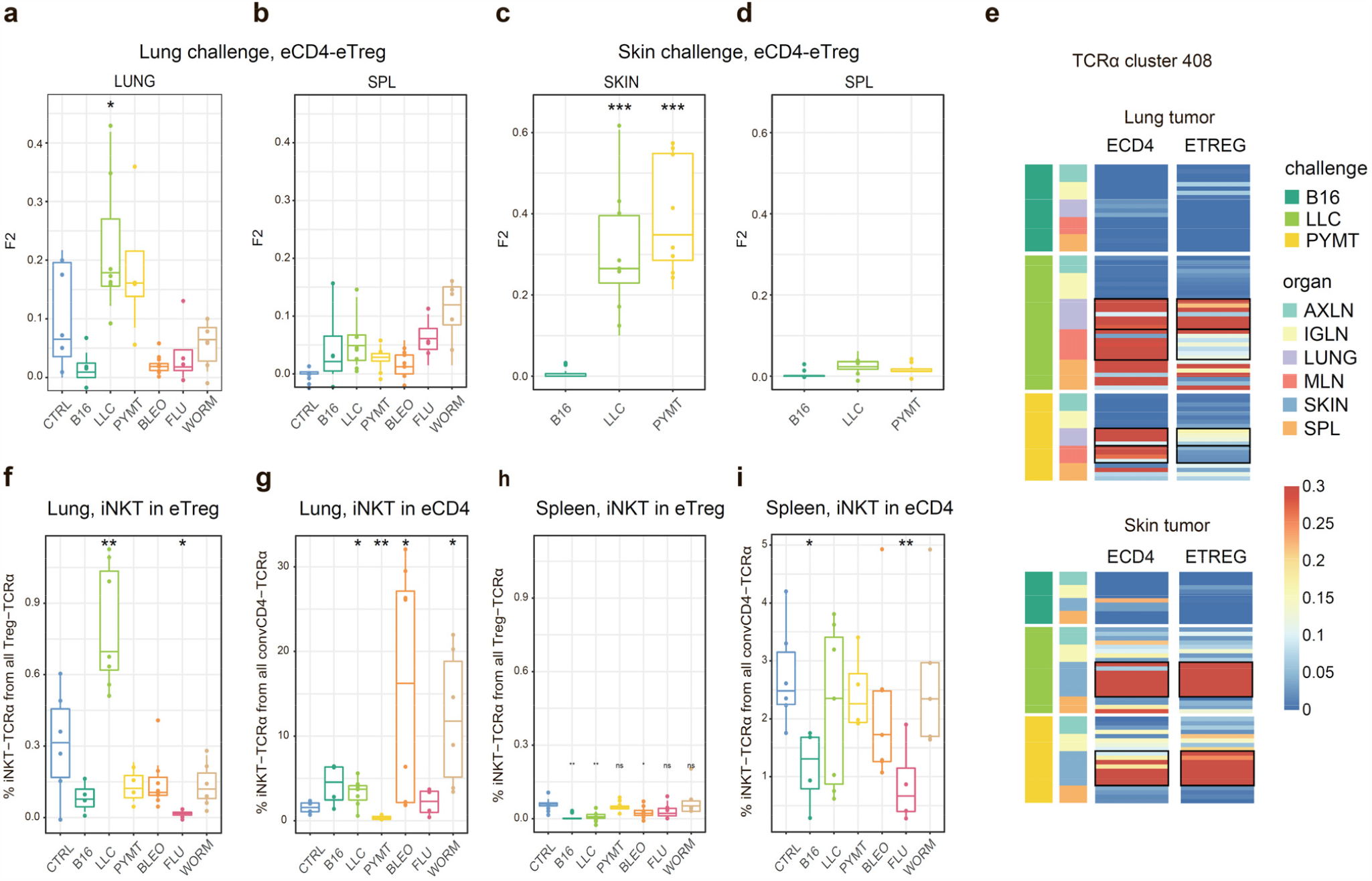
Estimated eCD4 to eTreg conversion. (**a–d**) Relative overlap between eTreg and eCD4 nucleotide-defined TCRα CDR3 repertoires upon lung (**a, b**) and skin (**c, d**) challenges. Overlaps are shown for the same organ (**a, c**) and spleen (**c, d**). Data were normalized to the 100 most frequent clonotypes and overlap was weighted by clonotype frequency (F2 metric). Clonotypes were matched on the basis of identical TRAV and TRAJ gene segments and identical TCRα CDR3 nucleotide sequences. Overlap was analyzed separately for each challenge for eTreg and eCD4 subsets obtained from the same mice. The closer two samples are, the higher the F2 metric, which reflects the overall frequency of shared clonotypes. **e**. Distribution of TCR cluster 408 in eTreg and eCD4 repertoires upon tumor challenge in lung and skin. (**f**) Treg-iNKT proportion of eTreg cells in lung upon lung challenge. (**g**) iNKT proportion of eCD4 cells in the lung upon lung challenge. (**h**) Treg-iNKT proportion of eTreg cells in spleen upon lung challenge. (**i**) iNKT proportion of eCD4 cells in the spleen upon lung challenge. P-values shown as: ^*^p < 0.05, ^**^p < 0.01, ^***^p < 0.001, and ^****^p < 0.0001 based on parametric t-test for each group versus control.

Foxp3^+^ Treg-iNKT cells do not naturally arise during development in the thymus, but can be induced and acquire suppressive functions in the periphery under particular pathophysiological conditions^26,27^. We specifically observed an increased proportion of Treg-iNKT cells (defined as classic TRAV11-CVVGDRGSALGRLHF-TRAJ18 TCRα) among all eTreg cells upon LLC challenge in lungs (**Fig. 4f**). This was not due to increased presence of eCD4 iNKT cells (**Fig. 4g**), suggesting induced iNKT conversion upon LLC lung challenge. The conversion, which was similar to the general conversion shown in **Figure 4a–d**, was local and was not observed systemically (*e*.*g*., in the spleen; **Fig. 4h**).

### Local lymphatic tissue-resident eTreg and eCD4 cells

Global TCRα CDR3 cluster analysis revealed that characteristic eTreg TCR motifs were present in distinct lymphatic tissues, including spleen and thymus, irrespective of the applied challenge (**Supplementary Figs. 1,2, Supplementary Tables 4**,**5**). To better illustrate this phenomenon, we performed MDS analysis of TCRα CDR3 repertoires for distinct lymphatic tissues, excluding the lungs due to their otherwise dominant response to the current challenge. This analysis demonstrated close proximity of eTreg repertoires obtained from the same lymphatic tissues upon all lung challenges and across all animals (**Fig. 5a, b**). These results indicate that distinct antigenic specificities are generally characteristic for eTreg cells that preferentially reside in particular lymphatic niches. Notably, the convergence of lymphatic tissue-resident TCR repertoires was less prominent for the eCD4 T cells (**Fig. 5c, d**).

**Figure 5.**
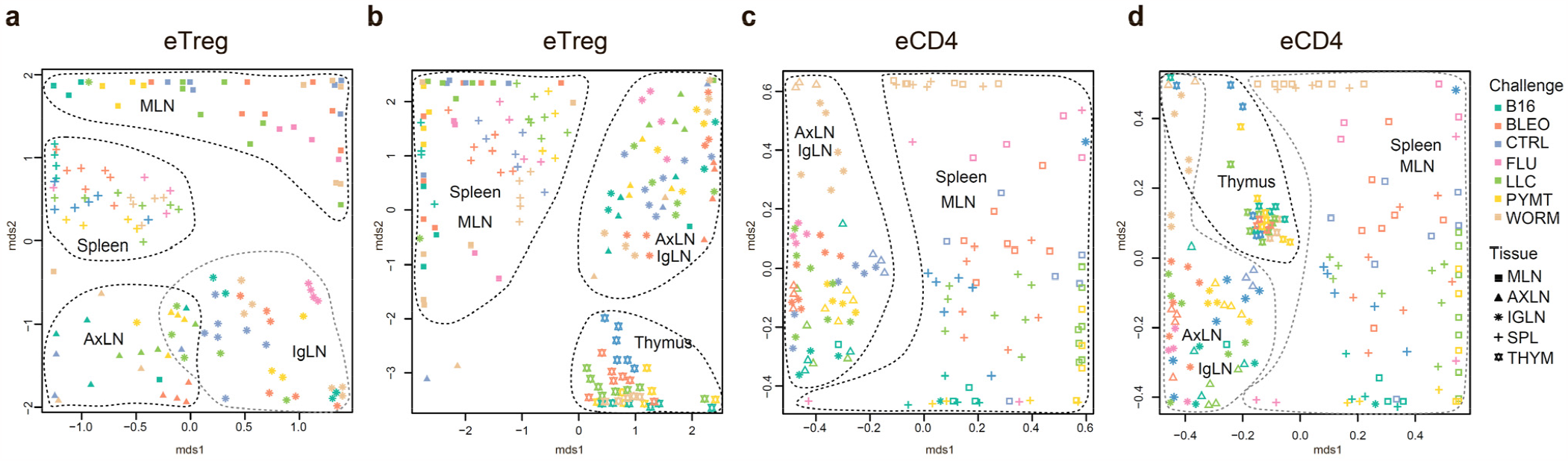
Lymphatic tissue-resident eTreg and eCD4 cells. (**a–d**) MDS plots showing relative overlap between amino acid-defined eTreg TCRα CDR3 repertoires as visualized using VDJtools. Repertoire overlaps are shown for (**a**) eTreg in MLN, spleen, AxLN, and IgLN; (**b**) eTreg in MLN, spleen, AxLN, IgLN, and thymus; (**c**) eCD4 in MLN, spleen, AxLN, and IgLN; and (**d**) eCD4 in MLN, spleen, AxLN, IgLN, and thymus. Data were normalized to the top 1,000 most frequent clonotypes and weighted by clonotype frequency (F2 metric). Clonotypes were matched on the basis of identical TRAV gene segments and identical TCRα CDR3 sequences. Euclidean distance between points reflects the distance between repertoires; the closer two samples are, the higher is the overall frequency of shared clonotypes. Mice were analyzed in an all-versus-all fashion, irrespective of whether the tissues were obtained from the same or distinct mice. Node colors correspond to the different challenges. Squares: MLN. Triangles: AXLN. Snowflakes: IGLN. Crosses: spleen. Stars: thymus.

## Discussion

We observed a prominent local clonal response to different antigenic challenges within the lung. Remarkably, this effect was comparable for both lung-infiltrating eCD4 and eTreg cells, suggesting similar amplitude for the effector and regulatory response (**Fig. 1a–c**). The TCR repertoires of the eTreg and eCD4 subsets were highly distinct across antigenic challenges (**Fig. 1d, e**). At the same time, eTreg repertoires were highly similar across mice for each challenge (**Fig. 2a, b**), and common TCRα CDR3 motifs dominated each response (**Supplementary Fig. 1**). That being said, the eTreg repertoires were similar for the three different cancer challenges, suggesting a response to shared tumor-associated antigens. Antigen-specific Treg response in lungs was reflected in the lung-draining MLN, but not in distant tissues including the AXLN, IGLN, and spleen (**Fig. 2b, Supplementary Fig. 1**), showing the local character of the antigen-specific regulatory response.

For the LLC and PYMT tumors—but not other antigenic challenges—the overall repertoires and dominant CDR3 motifs were shared between the two subsets, suggesting induction of eCD4-eTreg plasticity. Further analysis showed that eCD4 to eTreg conversion was strongest in the skin and lung following LLC and PYMT tumor challenge, and was observed in the site of challenge but not systemically (**Fig. 4a–e**). The presence of an increased proportion of Treg-iNKT cells among the total pool of eTreg cells in the lung upon LLC challenge provided further support for the notion of tumor-induced plasticity (**Fig. 4f–h**). This observation is of particular interest in the context of recent reports on innate-like T cell anti-tumor response^28^.

The local eTreg response to a given tumor type was clonally very similar upon lung and skin challenge, showing that the antigen-specific character of the eTreg response dominates over the tissue location of the initial challenge (**Fig. 3**).

Strikingly, we observed commonality between eTreg repertoires obtained from the same lymphoid tissue in different mice irrespectively of the antigenic challenge (**Fig. 5a,b**). This observation indicates that distinct antigenic specificities are characteristic for Treg cells with distinct tissue residence.

The eCD4 subset generally exhibited similar patterns to the eTregs, although the extent of convergence among eTreg repertoires was higher in most scenarios compared to the eCD4 subset, indicating the highly antigen-specific nature of the Treg response.

Altogether, our data demonstrate highly antigen-specific and distinct Teff and Treg responses to different challenges and support the use of the fixed TCRβ chain FoxP3-GFP mouse as a “digital” model of TCR response that benefits from access to highly convergent and easy-to-track TCRα CDR3 clusters.

## Methods

### Mice and challenges

C57BL/6J DO11.10 TCRβ transgenic mice (kindly provided by Philippa Marrack) and crossed to C57BL/6J Foxp3^eGFP^ TCRa^-/-^ mice. Age-matched DO11.10 TCRβ Foxp3^eGFP^ TCRa^+/-^ males were used for experiments. The animals were kept under specific pathogen-free conditions and studied at 6–9 weeks of age.

PyMT cell line was isolated previously from MMTV-PyMT tumor-bearing mice^29^, Lewis Lung Carcinoma (LLC) cell line was a gift of J.Massague (Memorial Sloan-Kettering Cancer Center) and B16-F10 melanoma cells were a gift from C. Ariyan (Memorial Sloan Kettering Cancer Center). All cell lines were grown in DMEM supplemented with 10% FBS, 1% l-glutamine, 1% penicillin-streptomycin, and 10 mM Hepes. Tumor lines were trypsinized and washed with serum-free DMEM. A dose of 1x 10^5^ cells in 200 uL DMEM was injected to each mouse intravenously (i.v) via the tail vein or subcutaneously (s.c) in right flank to induce lung or skin tumors, respectively. Influenza A/Puerto Rico/8/34 (PR8) virus was grown in the cavity of 10-day embryonated chicken eggs, and a dose 100 pfu in 35 uL PBS was used to challenge mice intranasally (i.n)^8^. *Nippostrongylus brasiliensis* L3 stage larvae were passaged and isolated from the feces of Wistar rats^30^. 500-700 infective L3 larvae were injected s.c to mice in 500 uL PBS. Bleomycin for injection was obtained from the pharmacy and each mouse was challenged i.n with a dose of 0.1 U in 35 uL PBS. Control mice were mock challenged with PBS. Antibodies for magnetic cell isolation and fluorescence activated cell sorting were purchased from Biolegend, Thermofisher and TONBO BIO. Mouse breeding, challenges and procedures were performed under protocol 08-10-023 approved by the Sloan Kettering Institute (SKI) Institutional Animal Care and Use Committee.

### Cell isolation, sorting of T cell subsets and library preparation

To identify and isolate tissue infiltrating T cells and exclude circulating blood lymphocytes, mice from different challenge groups were injected i.v with 0.5 ug of APC anti-CD45 antibody (clone 30-F11) 3 min before euthanasia^31^. Mice bearing lung and skin tumors were euthanized 2 weeks after challenge and influenza, *Nippostrongylus brasiliensis* and bleomycin experimental mice at day 9 after challenge.

Lungs from all challenge groups were flushed from excess blood via intracardiac injection with 10 mL of PBS, and each lobe was cut into small pieces and digested with Collagenase A (Roche, 1mg/mL), DNAse (30 ug/mL) in DMEM 2% FBS for 30 min at 37°C in an orbital shaker. Skin tumors were dissected, minced into small pieces and digested with Collagenase A as described above for lung samples. Single cell suspensions were spun and resuspended in sterile EDTA containing FACS buffer (PBS 2% FBS, 1mM EDTA). Spleens and individual lymph nodes were dissociated with frosted glass slides into single-cell suspensions in FACS buffer. All cell suspensions were filtered with 70 uM strainers (BD Biosciences) and kept on ice until further use.

Individual cell suspensions were enriched for CD4^+^ T cells using a custom cocktail of biotinylated antibodies (F4/80(clone BM8), anti-mouse I-A/I-E (clone M5/114.15.2), anti-mouse B220(clone RA3-6B2), anti-CD11b (clone M1/70), anti-mouse gamma delta TCR (clone GL3), anti-mouse CD8a (clone 53-6.7), anti-mouse CD11c (clone HL3), anti-mouse Ter119 (clone Ter119), anti-mouse Ly6-G (clone 1A8) at a concentration of 10 ug/mL for each antibody) followed by negative selection with LS columns (Miltenyi). Negative fractions were stained with anti-CD4 (GK1.5, BV605), CD8b (YTS157.7.7, AF700), CD44 (IM7, EF450), CD62L (MEL-14, PE-TexasRed), CD90.2 (clone 30-H12, APC-Cy7), CD45 (clone 30-F11, BV510) and Vb8.2 (clone KJ16-133, PE) antibodies. Effector Treg cells and effector CD4^+^ T cell subsets were sorted using an Aria-II Cell Sorter (BD Biosciences). Briefly, total effector CD4^+^ T cells were gated as Vb8.2^+^ CD4^+^ CD44^hi^ CD62L^+^ CD90^+^ CD45-BV510+ CD45 APC-cells. Thereafter, from this gate, conventional effector CD4^+^ T cells as CD44^hi^ CD62L^-^ Foxp3-GFP^-^ cells and effector Treg cells as CD44^hi^ CD62L^lo^ Foxp3-EGFP^+^ were sorted individually into separate tubes, spun, and lysed in buffer RLT plus (Qiagen) and froze at -80°C until further use. RNA purification from individual samples (RNeasy Micro Kit, Qiagen), TCR library preparation from cDNA and Library Next Generation Sequencing has been previously described^18^.

### TCR repertoire extraction

Raw 150+150 nt sequence data were analyzed using MIGEC software version 1.2.7. UMI sequences were extracted from demultiplexed data using the Checkout utility. Then, the data were assembled using the erroneous UMI filtering option in the Assemble utility. The minimum required number of reads per UMI was set at two for most tasks. In-frame TCRα and TCRβ repertoires were extracted using MiXCR software (version 3.0.13). Normalization, data transformation, in-depth analyses, and statistical calculations were performed using VDJtools software version 1.2.1^24^. R scripts were used to build figures.

### TCRα repertoires diversity analysis

TCRα repertoire diversity was assessed using several widely used metrics, including observed diversity, normalized Shannon Wiener index, and Chao1. For normalization, all diversity metrics were obtained for datasets that had been downsampled to 1,000 randomly-chosen, UMI-labeled TCRα cDNA molecules. Samples with UMI < 700 were excluded from analysis.

### TCRα repertoires overlap analysis

Repertoire overlap was analyzed using the weighted F2 (reflecting the proportion of shared T cells between paired repertoires) metric in VDJtools software version 1.2.1. For amino acid overlap metrics calculations, we selected top 1,000 largest clonotypes from each cloneset. Samples with clonotype counts <700 were excluded from analysis. The top N clonotypes were selected as the top N clonotypes after randomly shuffling the sequences and aligning them in descending order. This was done in order to get rid of the alphabetical order for clonotypes with equal counts (e.g. count = 1 or 2).

### eCD4-to-eTreg conversion analysis

The top 100 clonotypes were extracted from all samples. F2 (weighted on clonotype size) overlaps in terms of nucleotide CDR3 sequence plus identical V and J were calculated for matched pairs of samples as shown in **Supplementary Table 3**. Samples with clonotype counts <100 were excluded from analysis. The top N clonotypes were selected as the top N clonotypes after randomly shuffling the sequences and aligning them in descending order. This was done in order to get rid of the alphabetical order for clonotypes with equal counts (e.g. count = 1 or 2).

### Statistical analysis

Results are shown as mean ± SEM. Statistical analyses were performed on processed datasets in R. Multiple parameter inferences were estimated using parametric t-test.

### TCRα clusters

TCRα sequence homology clusters were generated using the TCRNET algorithm^32^. All TCRα CDR3 sequences were pooled, and unique sequences were used to build a graph. Edges of this graph connected CDR3s that differed by a single amino acid mismatch. CDR3s having more neighbors than expected by chance were selected, and connected components from selected CDR3 sequences were used as clusters. As a baseline, we used random mouse TCRα VDJ rearrangements. Each cluster was assigned a frequency based on the total frequency of T cells encoding corresponding CDR3s within a sample. The CDR3 cluster frequency matrix was then subjected to hierarchical clustering using the ‘aheatmap’ R package with default parameters.

## Acknowledgments

Authors are grateful to Alexander Y. Rudensky for his invaluable contribution to the work in all aspects, and to Michael Eisenstein for the helpful edits. Supported by the grant from the Ministry of Science and Higher Education of the Russian Federation 075-15-2019-1789.

## Data Availability

All repertoire data used in the manuscript are available on figshare: https://figshare.com/articles/dataset/Convergence_plasticity_and_tissue_residence_of_regulatory_and_effector_T_cell_response/22226155

---

**Supplementary Fig. 1.**
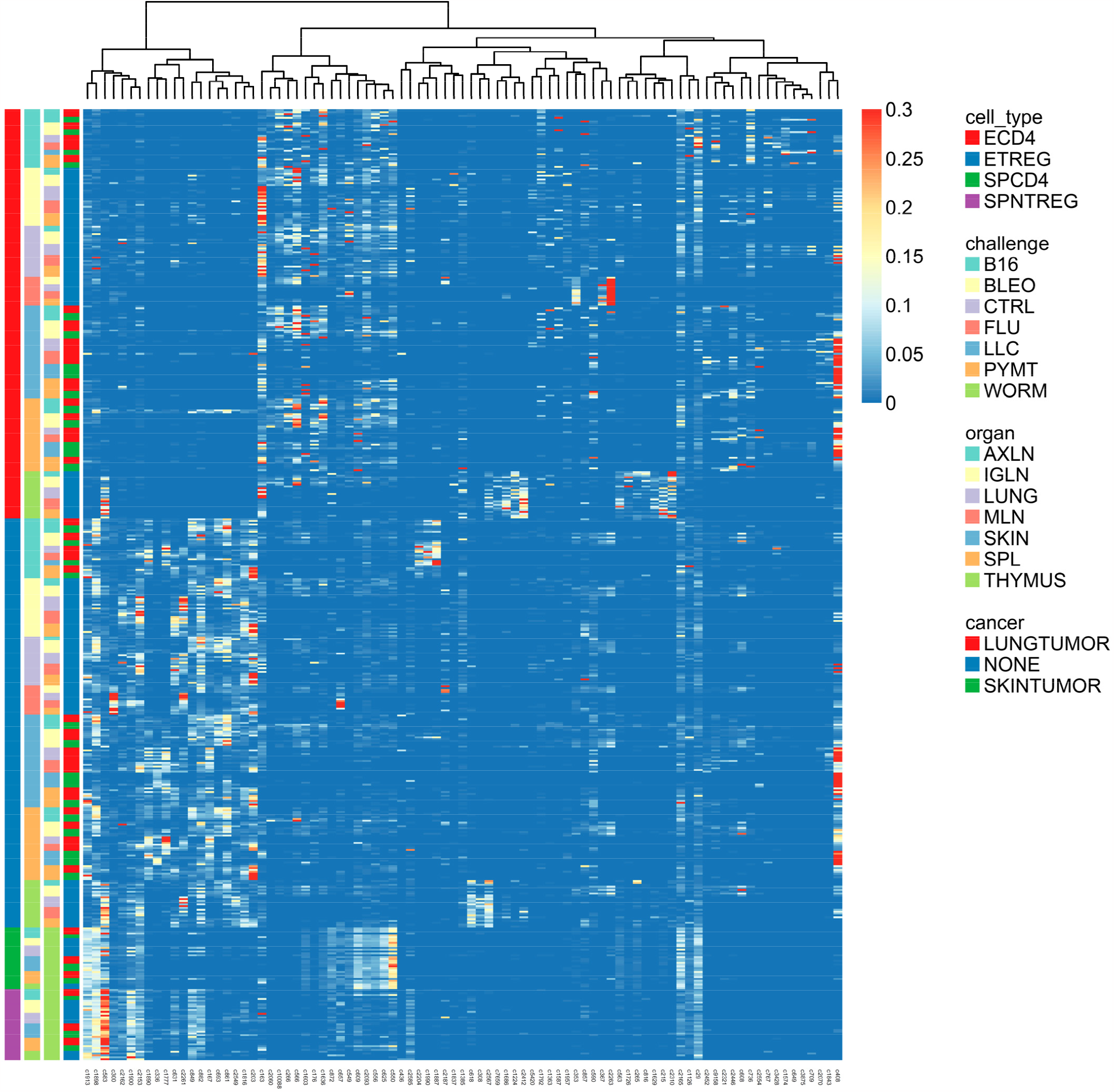
TCR group behavior in response to different antigenic challenges. In this TCR homology cluster expression matrix, each column corresponds to a TCR sequence homology cluster (*i*.*e*., group of TCR clonotypes having similar TCRα CDR3 sequences) and each row corresponds to a particular condition (*e*.*g*., tissue type, antigenic challenge, etc). The color of each cell highlights the overall (sum of frequency across all clonotypes) expression of each cluster in a given condition. Rows are sorted by cell type, challenge, organ and cancer type as highlighted by the colored bands at left; columns are sorted according to hierarchical clustering of TCR cluster expression.

**Supplementary Fig. 2.**
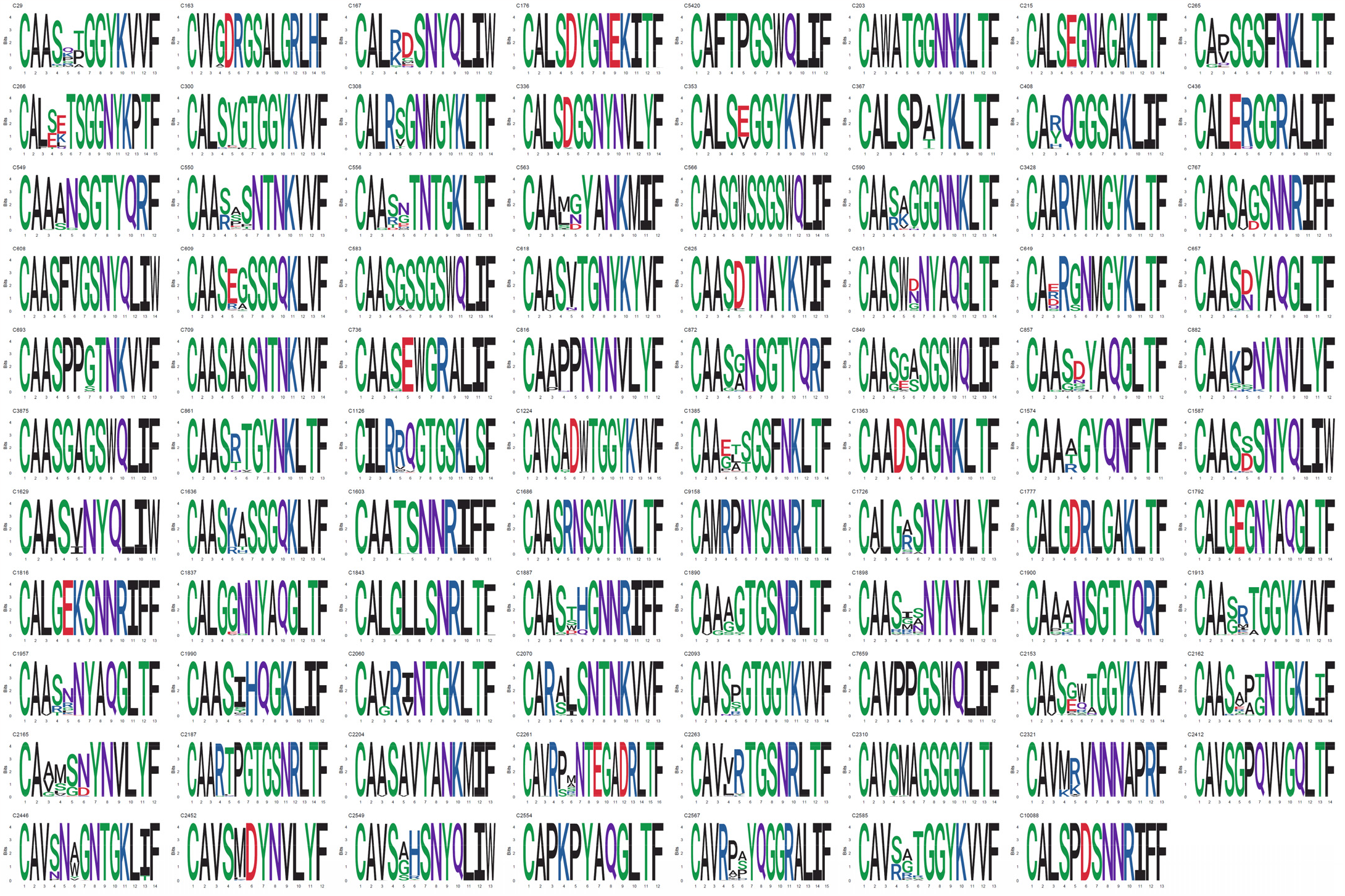
All TCR clusters.

