## Supplementary figures and images for "Convergence, plasticity, and tissue residence of regulatory T cell response via TCR repertoire prism"

### Supplementary Figure 1

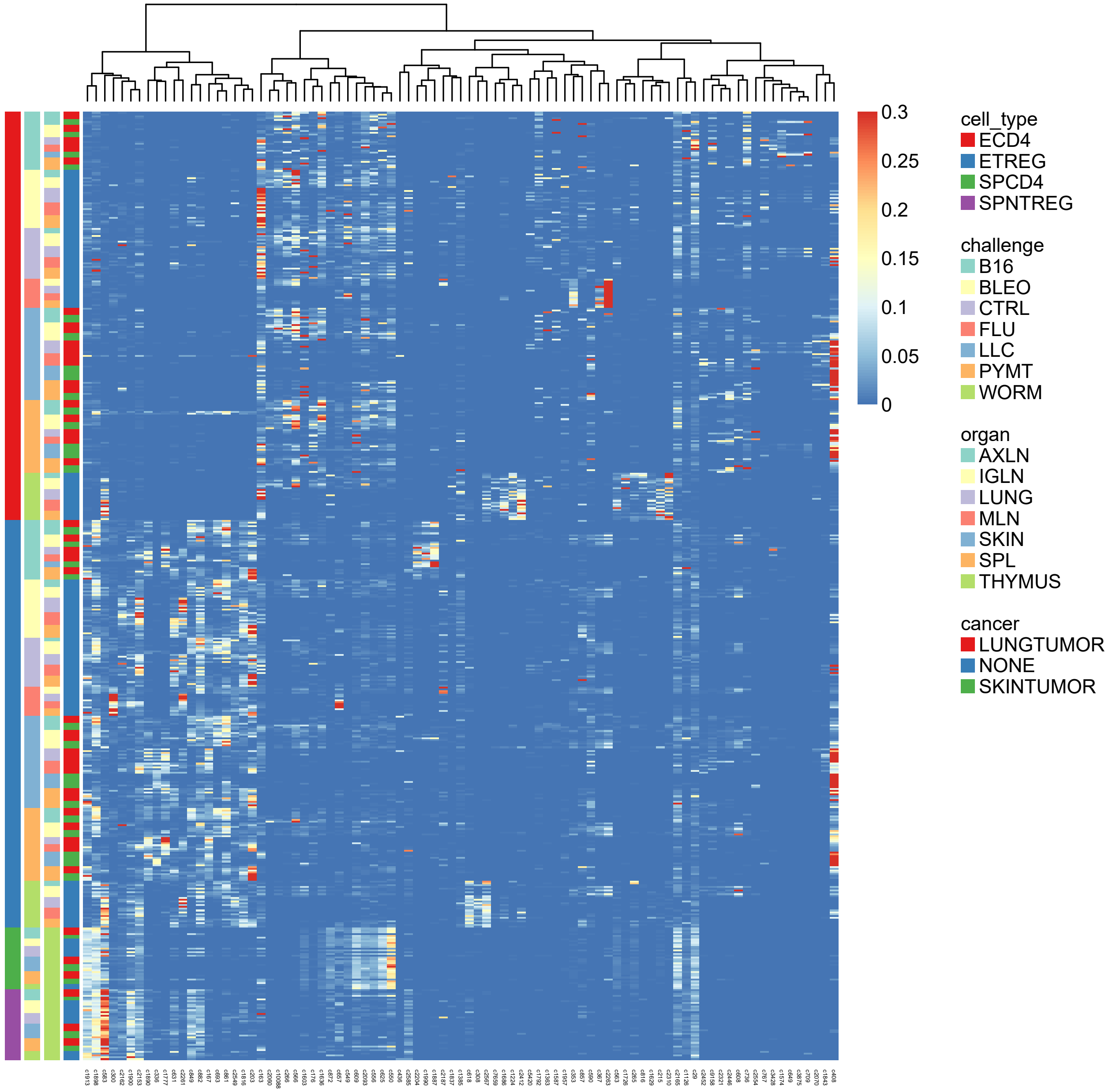

### Supplementary Figure 2

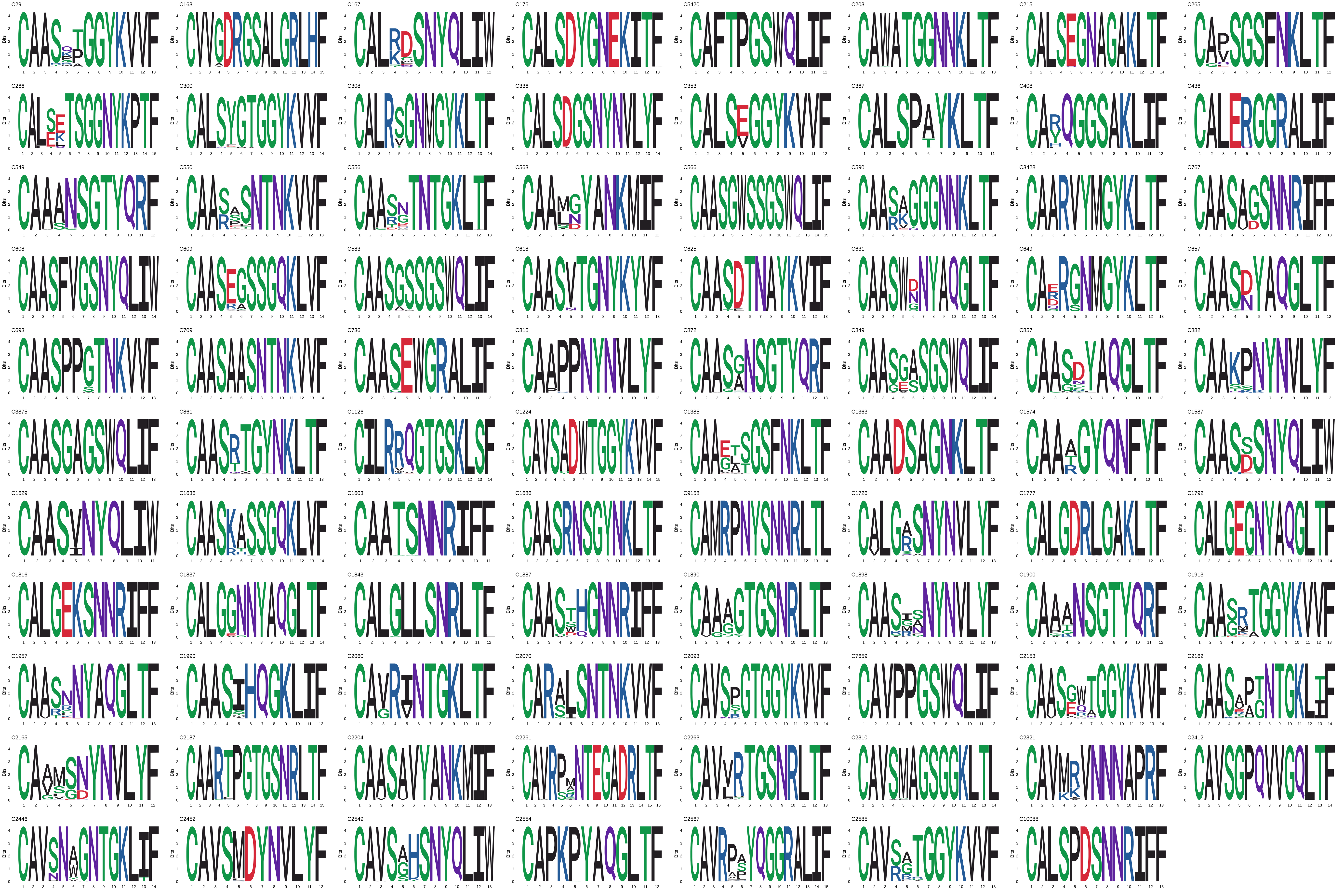
